# Modeling gene expression networks to predict interchromosomal organization during human embryonic stem cell differentiation

**DOI:** 10.1101/318899

**Authors:** Kyle V. Laster, Arturo G. Garza-Gongora, Elizabeth Daley, Alexey Terskikh, Evangelos Kiskinis, Erica D. Smith, Steven T. Kosak

## Abstract

Cellular differentiation occurs through the regulation of lineage-specific gene expression networks that are facilitated by the spatial organization of the genome. Although techniques based on the chromatin conformation capture (3C) approach have yielded intrachromosomal genome-wide interaction maps, strategies to identify non-random interchromosomal associations is lacking. Therefore, we modeled the genomic organization of chromosomes based on the regulatory networks involved in the differentiation of pluripotent human embryonic stem cells (hESCs) to committed neuronal precursor cells (cNPCs). Importantly, transcriptional regulation has been demonstrated to be a driving force in non-random genome organization. Thus, we constructed coarse-grained *in silico* networks using gene expression data to identify potential physical associations among chromosomes occurring *in situ* and then analyzed the three-dimensional (3D) distribution of these chromosomes, assessing how their associations contribute to nuclear organization. Our analysis suggests that coordinate regulation of differentially expressed genes is correlated with the 3D organization of chromosomes in hESC nuclei induced to differentiate to cNPCs.

**Author Summary:** The cellular commitment and differentiation of stem cells is a hallmark of metazoan development. The ultimate fate of a stem cell is defined by the synergistic modulation of key gene regulatory networks within the nucleus. In our work, we formulate an *in silico* model describing how the similarity in the expression profile of differentially regulated gene networks is correlated with the higher-order organization of chromosomes during differentiation from human embryonic stem cells (hESCs) to committed neuronal precursor cells (cNPCs). Using graph statistics, we observe that the genome networks generated using the *in silico* model exhibit properties similar to real-world networks. In addition to modeling how gene expression relates to dynamic changes in chromosome organization, we test the model by calculating the relative proximity of multiple chromosome pairs using 3D fluorescence *in situ* hybridization (FISH). While various chromosomal properties, including gene density and overall length, have been attributed to chromosome organization, our previous work has identified the emergence of cell-type specific chromosomal topologies related to coordinate gene regulation during cellular differentiation. Here we extend these findings by determining whether our *in silico* model can predict chromosome association based upon coordinate gene expression. Our work supports the idea that gene co-regulation, in addition to inherent organizational constraints of the nucleus, influences three-dimensional chromosome organization.

## Introduction

The vertebrate genome exhibits patterns of non-random organization during cellular differentiation, with developmentally regulated genes positioned within the nucleus according to their expression [1,2]. The periphery of the nucleus and its interior are compartments that have been demonstrated to be transcriptionally repressive and permissive, respectively [3]. Co-regulated genes have also been shown to cluster according to their expression status and to spatially associate with nuclear bodies (NBs), such as nuclear speckles and promyelocytic leukemia (PML) bodies [4-7]. Moreover, the chromosomes on which co-regulated genes are encoded also demonstrate non-random organization within the nucleus [8].

Seminal studies indicate that chromosomes can be organized according to gene density or size, with gene rich or smaller chromosomes occupying more centralized positions within the nucleus and gene poor or larger chromosomes at the periphery [9-11]. Inherent organizational constraints also appear to facilitate associations of chromosomes through intermediate structures, such as the nucleolus and the nuclear lamina [12-14]. Additionally, patterns of non-random chromosomal association are evident among different cell types and cellular states [15-17]. Importantly, we have previously reported that chromosomes enriched for genes co-regulated during murine erythropoiesis demonstrate both homologous and heterologous interactions that correlate with lineage-specific expression profiles [18, 19]. Aberrations in the coordination of gene expression and genomic organization also reveal deleterious effects in development and disease [20-22].

The three-dimensional (3D) structure of eukaryotic chromatin also reveals non-random organization. Advances in chromatin conformation capture (3C) derived methodologies (in particular Hi-C), which identify transiently interacting chromatin through sequencing of *in situ* cross-linked DNA, have shown that chromatin is organized into topologically associated domains (TADs) of ~1Mb in humans [23-25]. TADs are flanked by CTCF-binding sequences that appear to constrain the interactivity of the chromatin confined within a given region [26, 27]. The size and distribution of TADs appear to be cell-type invariant; however, the genes contained within each distinct TAD boundary have been found to be coordinately regulated and may inform the 3D nuclear localization of TADs [28, 29]. Although 3C based techniques have been successful in providing genome wide resolution of *cis*-interacting chromatin, they have yet to fully characterize interchromosomal associations, as these interactions are hard to detect by these strategies.

Pluripotent human embryonic stem cells (hESCs) give rise to the three germ layers that generate the organism [30]. Pluripotency is maintained via a core transcriptional network governed by robust expression of Oct4, Sox2, and Nanog. This regulatory network influences cellular identity in three interconnected ways: preservation of chromatin plasticity, repression of key genes involved in lineage commitment, and modulating genomic architecture. Increased chromatin plasticity occurs due to reduced concentrations of heterochromatin and chromatin stabilizing proteins such as the nuclear lamina protein lamin A/C [31-34]. As a result chromatin dynamics of the pluripotent genome are enhanced, permitting increased interaction of distal regulatory elements with target sequences in a developmentally regulated manner [35]. This unique euchromatic conformation also contains an epigenetic signature that is permissive to basal-level expression of the hESC genome [36], which is suggested to minimize expression of developmentally regulated genes while simultaneously priming them for activation in the event of an extracellular commitment cue. Upon commitment, the diminution of the master transcriptional regulators of pluripotency results in extensive changes in the chromatin landscape, thereby influencing gene expression toward lineage restricted pathways [37-39]. Thus, hESC differentiation is an ideal model to explore the interplay between interchromosomal association and gene expression networks.

Here we test whether we can predict, based on the transcriptional profiles of developmentally regulated genes, 3D interchromosomal proximity within interphase nuclei in hESCs (H9) during their differentiation into committed neuronal progenitor cells (cNPCs) [40]. To determine gene loci that may synergistically influence interchromosomal associations, we identified differentially expressed genes based on microarray data from a 10-day differentiation time course. Next, we aligned each gene of the enriched gene set along its encoding chromosome and performed a pairwise comparison of their normalized expression scores using a non-Euclidean distance metric. We constructed *in silico* networks that rank potential *in situ* associations between chromosomes based on the pairwise comparison of their gene expressions at each point of the differentiation time course. To test if our *in silico* predictions faithfully recapitulate *in situ* genome organization, we assessed metrics of chromosome association at three time points of differentiation using 3D fluorescence *in situ* hybridization (FISH) analysis with whole chromosome paints. Our results demonstrate that chromosomes are distributed non-randomly within interphase nuclei of differentiating hESCs and that similarity in expression of co-regulated genes is useful in predicting trends in physical proximity of heterologous chromosomes. We believe this approach will be an important tool for identifying cell-type specific genome topologies in differentiation and disease.

## Results

### hESC differentiation to cNPCs reveals dynamic changes in expression

As a first step in our analysis, we determined the co-regulatory gene set by conducting Significance Analysis of Microarrays [41], on a quantile normalized dataset of global changes in gene expression upon commitment of pluripotent H9 ESCs to cNPCs over a 10-day differentiation time course [42]. Using a False Discovery Rate (FDR) of 13%, we found that 2040 out of 19284 represented genes demonstrate differential expressions: 1570 genes were up-regulated while 470 genes were down-regulated over the 10-day differentiation time course (S1 Table) and the genes were assigned to their chromosomal positions (please refer to reference 40 for elaboration of the genes identified). Linear regression analysis reveals that on a genome-wide level, the greater number of differentially expressed genes on a given chromosome has a strong correlation with its number of significant bins (or domains) (r=.629) (S2 Fig). These results indicate that the gene set driving differentiation of hESCs to cNPCs is non-randomly distributed along the linear genome. More importantly for our modeling, these data support our hypothesis that the similarity of expression profiles among heterologs may predict their nuclear proximity.

We next examined the dynamic behavior of gene expression changes occurring during the differentiation of hESCs to cNPCs in order to determine how the profile was modulated. A clustergram of the expression profile reveals grouping into two broad classes of up-regulated and down-regulated genes, but closer inspection of the heatmap reveals nuanced behavior within these clusters (Fig 1A). In order to visualize predominate classes of behavior within our system, we naturally partitioned the data into four classes of expression based on the L-method [42] before using *k*-means clustering. This particular algorithm determines ideal cluster number by minimizing the root mean squared error between two lines iteratively fit to the merge distance of a finite number (~100) of divisive hierarchical clusters. The *k*-means analysis reveals four distinct patterns of gene regulation induced upon the onset of neurogenesis (Fig 1B, S3 Table).

**Fig 1.**
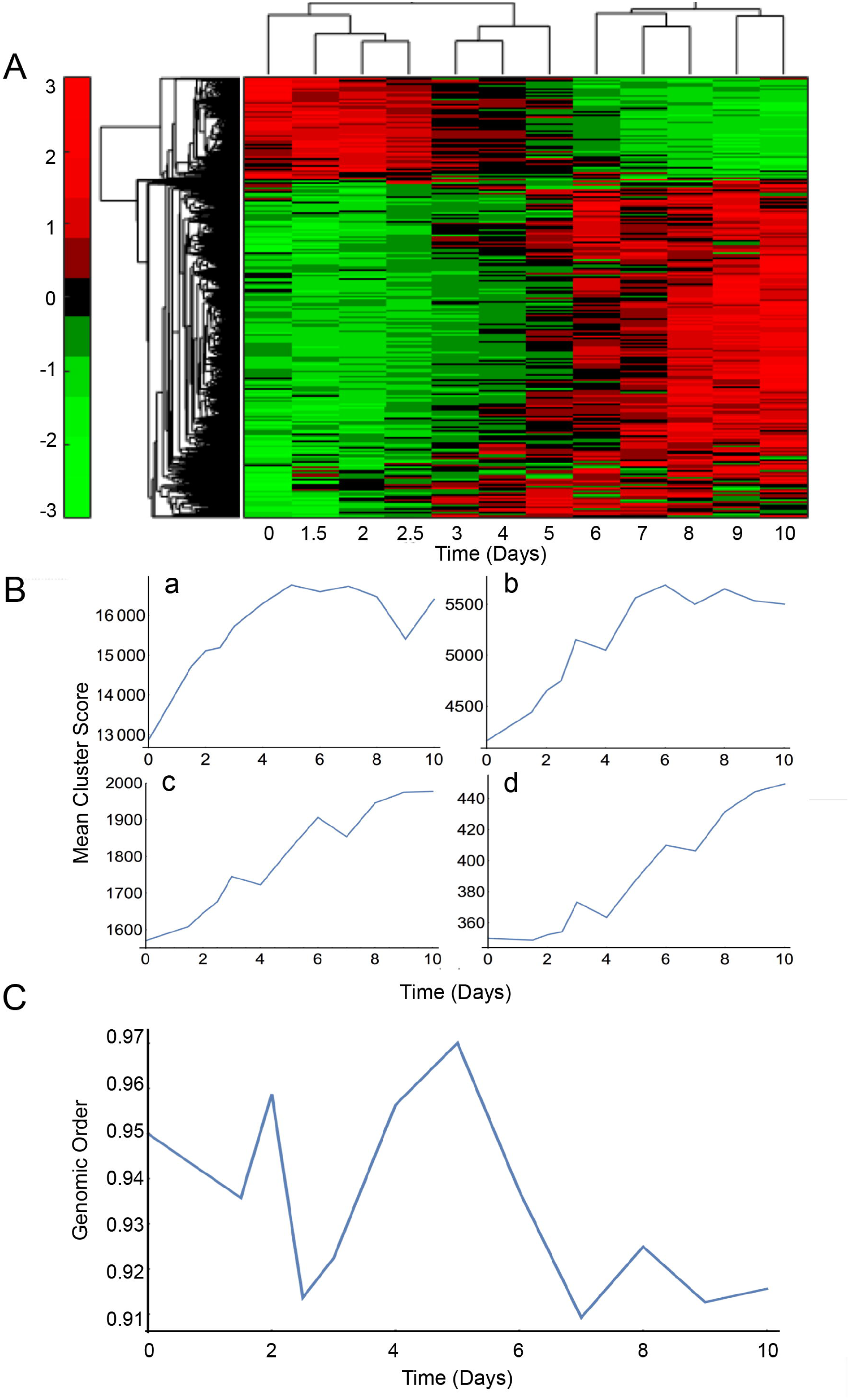
Gene expression is dynamically modulated during neuronal commitment. A: Clustergram generated using the Bioinformatics toolbox in MATLAB visualizing changes in expression of genes induced upon neurogenesis. The 2040 differentially expressed genes identified using Significance Analysis of Microarrays were hierarchically clustered based upon their normalized expression values over the 10-day differentiation time course. The expression profile is clustered into two broad classes of up-regulation (green to red) and down-regulation (red to green). Each time point is indicated below the clustergram. The legend detailing expression levels is shown to the left. B: *k*-means clustering profile detailing four main patterns of expression during neuronal commitment. Utilization of the L-method recommended partitioning each gene into one of four discreet classes (a-d) based upon its normalized expression score at each time point of the differentiation. Although each class has a distinct kinetic signature, there are some shared features in (b-d) that highlight behavioral synergy within the differentially expressed gene set. C: Genomic order is a quantitative metric that identifies the largest fraction of differentially expressed genes that are synergistically expressed. It is measured by calculating the number of genes within the giant component at each individual time point as a ratio to the total number of differentially expressed genes identified using Significance Analysis of Microarrays. A value of 1 indicates the highest degree of transcriptional synergy between genes of the differentially expressed set, with the opposite being true for values closest to 0. The genomic order of our identified gene set roughly behaves in a sinusoidal fashion, which follows important transitions during cNPC differentiation. The genomic order of our dataset reaches its apex at Day 5 suggesting that most genes are synergistically expressed towards the midpoint of the differentiation cycle during the neurosphere phase (See S7 Fig).

Our approach provides insight into how the absolute behavior of our co-regulated gene set changes over the differentiation time course, but it does not indicate relatedness in expression of the genes with respect to one another. To illustrate the dynamically evolving nature of global gene expression changes occurring over the differentiation time course, we hierarchically clustered the Symmetrized Kullback-Leibler (SKL) divergence of the normalized expression scores between all of our differentially expressed genes in a pair-wise fashion. The unbounded numerical output of the SKL ratio serves as a non-Euclidean distance metric between the gene expression scores of our lineage-specific gene set, with 0 being the closest possible distance between elements. Gene distribution among the clusters appears highly polarized, with the largest cluster comprising between ~91-97% of our differentially expressed set over the time course (Fig 1C). This clustering feature, known as the giant component [19], indicates the largest fraction of synergistically expressed genes driving the process of commitment at each individual time point. Importantly, with respect to our dataset, this illustrates that co-regulatory networks involved in the maintenance of pluripotency become destabilized upon initial commitment. As time proceeds, genes involved in the induction and maintenance of neurogenesis experience the greatest similarity in expression midway through the differentiation, before ultimately diverging from the component toward the end of the time course. Thus, gene expression order is reduced at the cNPC stage when the progenitors are poised for subsequent differentiation cascades (Fig 1C). These changes in genome order are not restricted to hESCs differentiating to cNPSs. We performed the same analysis on hESCs undergoing mesodermal differentiation and observed a similar pattern of sinusoidal oscillation over an 8-day time course (S4 Fig). These results imply a general underlying trend that occurs upon cell specification and differentiation.

### In silico networks predict preferential chromosome associations at various time points during differentiation

The results above suggest that a large fraction of our differentially expressed gene set are highly similar in expression, but does not indicate how expression similarity of these genes could potentially contribute to genome wide chromosome organization. To test our hypothesis of expression mediated interchromosomal interactions, we modeled the likelihood of specific chromosome associations by creating sparse networks based on expression similarity of differentially expressed genes, anchored upon their respective chromosomes. We used the mean pairwise SKL divergence of all genes between two chromosomes to serve as an indicator of potential *in situ* association, with the lowest values determining an increased likelihood of interaction (see Materials and Methods, Bioinformatics). The diploid genome functions as a complete network composed of 46 nodes (all of the chromosomes) and the 1035 edges that represent the probability of association between all nodes (Fig 2A). However, due to the inability to map in real time all RNA transcripts to their chromosome of origin, it is currently impossible to distinguish homolog contribution of gene expression; therefore, we modeled the haploid genome with 23 nodes.

**Fig 2.**
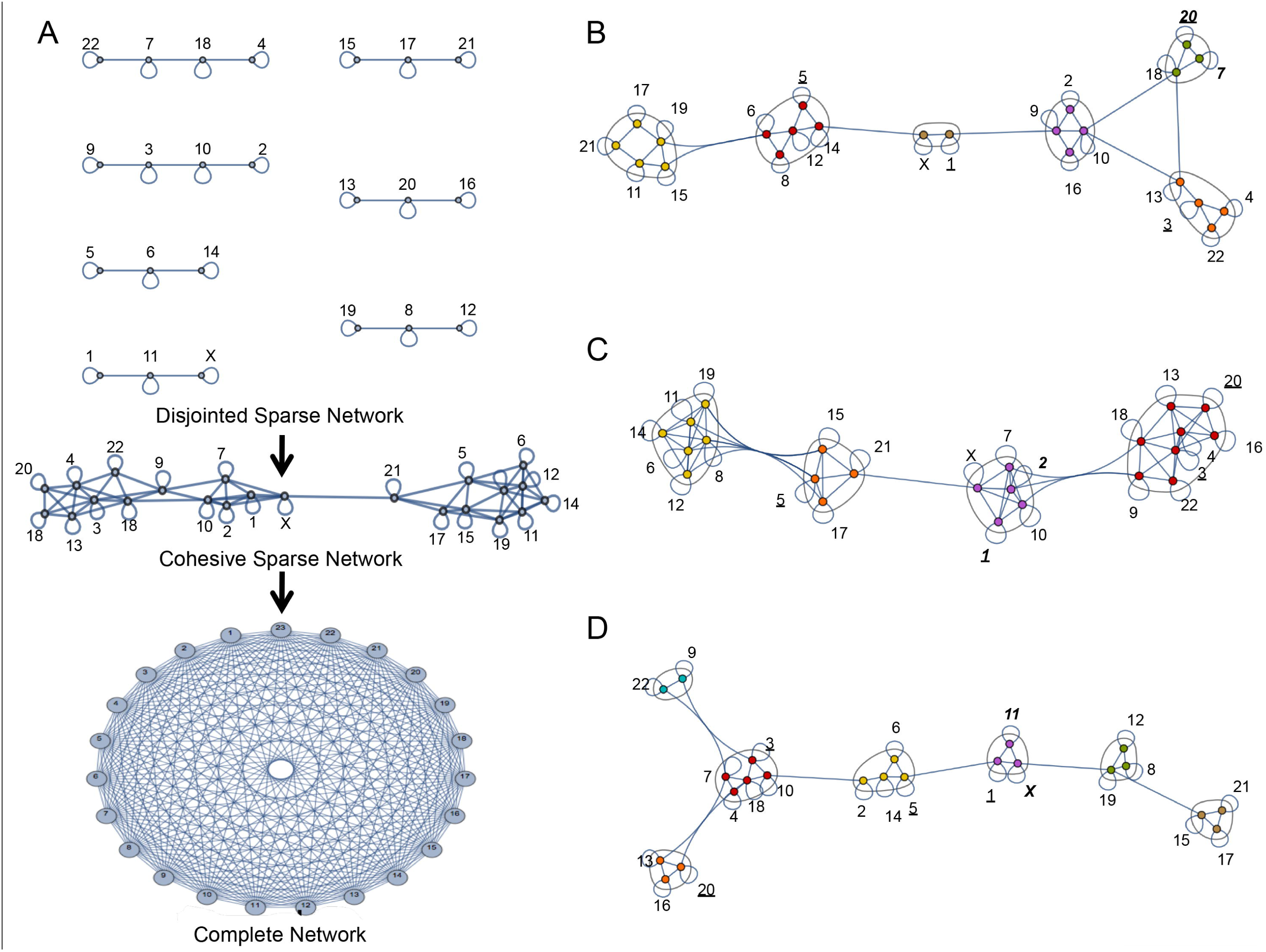
Developing *in silico* genome networks to predict expression based interchromosomal associations. A: Schematic detailing network formation from individual nodes. Node connectivity is contingent upon transcriptional similarity, as determined by the normalized mean SKL of the differentially expressed genes anchored upon their respective chromosomes. Early edge addition cycles result in the formation of disjointed sparse networks (top). Eventually, subsequent addition of edges results in the formation of the cohesive sparse network (center) and finally the complete network (bottom). The cohesive sparse network provides an emergent snapshot of the nodes that should be most likely to physically associate *in situ*. B-D: Network modularity illustrates potential *in situ* chromosome association. Cohesive graphs from Day 0 (B), Day 5 (C), and Day 10 (D) depict the formation of distinct graph neighborhoods based on the property of modularity. Member nodes of each color-coded, encircled neighborhood interact at a higher frequency with each other than with the nodes in any other neighborhood. Chromosomes chosen for *in situ validation* of our model are indicated in bold and italic (experimental) and underline (control).

We constructed haploid genome networks in a bottom-up fashion, utilizing a rank-based association schema, whereby directional edges were assigned between nodes (representing chromosomes) with the lowest mean SKL values (Fig 2A). Edges were iteratively added between all nodes until a unified graph was formed, with one edge per node added at each cycle. The number of edge cycles needed to achieve graph connectivity ranged between 2-4 addition cycles over the entire time course. Edges added during early iterations should indicate increased similarity in expression, and higher propensity for *in situ* association, than edges added during later iterations. As such, we hypothesized that these early edges would assist in guiding the genomic organization of chromosomes. Additionally, we predicted that bi-directionality between nodes occurring early in edge addition would indicate even higher propensity for *in situ* association due to increased transcriptional synergy between those nodes.

To visualize potential associations among the chromosomes within the network during the differentiation of hESCs to cNPCs, we created network communities based on the property of modularity at days 0, 5, and 10 that reveal discreet node neighborhoods highly enriched with connections to one another (Fig 2B-D). These modularity-based neighborhoods illustrate increased clustering among community members as opposed to the rest of the network, which we hypothesize to be indicative of reduced proximity amongst their representative chromosomes within the nuclear compartment due to the transcriptional coordination of our lineage-specific gene set. As the number of edges used in the construction of the cohesive network may influence the number of nodes within each neighborhood, due to increased probability of node association, we quantified the linkage density of the networks at each time point. As expected, there is a correlation between the network linkage density, a metric of overall node connectivity, and the number of neighborhoods at each time point in the network because of an overall increased edge number (S5A Fig).

Comparing topological metrics between networks can be used to assess their similarity to one another. Common metrics that may be used in such comparisons are degree distribution, the global clustering coefficient (GCC), and network assortativity [43]. Degree distribution describes edge number with respect to all nodes. The GCC describes the degree to which nodes within the network cluster together. Network assortativity is a quantitative topological metric spanning from −1 to 1 that reveals trends in network polarity; negative and positive assortativity indicate relationships between nodes of a dissimilar degree and a similar degree, respectively. Two meaningful networks that can serve as controls to our expression-based chromosomal interaction (genome) network are the Erdős-Rényi based random network and the Barabási-Albert based scale-free network, both created with the node-edge parameters we have utilized in our model. Several well-studied networks, such as the Internet and various protein-protein interaction matrices [44, 45], have been shown to be scale-free, possessing a degree distribution following the power law. By contrast, the random network does not adequately model any biological or real-world networks [46]. Therefore, we compared several measurements of network topology between our emergent genome network, the Erdős-Rényi random network, and the Barabási-Albert scale-free network. Additionally, we assessed how stable the topological metrics were by iteratively rewiring the genome network in a random fashion (s and l; 5 and 10 rewiring events, respectively).

We first compared degree distribution of the emergent genome network to the normal Erdős-Rényi random network at all time points (S5B Fig). Whereas the random graphs have symmetrical degree distributions at each point of the differentiation time course, the degree distribution of our *in silico* chromosome association networks tend to have positively skewed distributions, indicating a departure from normality (S5B Fig). This skewed distribution illustrates that there are many chromosomes that have few connections and a few chromosomes that have many connections; this observation is reminiscent of the power law distribution observed with scale-free Barabási-Albert networks. Moreover, this result suggests that our *in silico* chromosome association genome network possesses properties similar to real-world networks.

To assess the degree to which the nodes cluster within the emergent genome network, we compared its GCC to the rewired genome networks and the Erdős-Rényi random and Barabási-Albert scale-free control networks (Materials and Methods, Bioinformatics) (Fig 3A and S6A Fig). The nodes of many real-world networks tend to form highly clustered groups possessing dense edge concentrations relative to random networks. As indicated above, the degree distribution of the genome network exhibits a significant departure from normality; thus, we hypothesized that a heightened GCC would reflect this property when compared to the control networks. Although the behavior of all three networks positively correlates to linkage density, we find that the GCC of the genome network is elevated at all time points when compared to controls (S6A Fig), thereby supporting our hypothesis. As the genome network is rewired, the connectivity is progressively compromised in magnitude with the 5-edge and 10-edge rewirings (Fig 3A).

**Fig 3.**
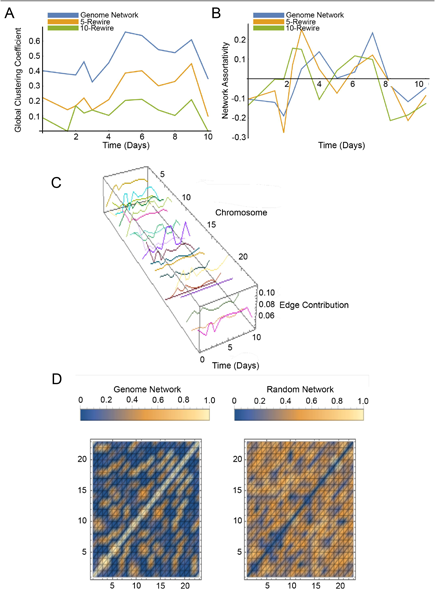
Cohesive genome networks exhibit unique topological organization that collapses upon perturbation. A: The global clustering coefficient (GCC), a metric of detailing the extent of network connectivity, is assessed for the genome network (blue), a 5-edge rewiring (yellow), and a 10-edge rewiring (green). The graph illustrates that the unperturbed genome network exhibits the highest degree of connectivity at all time points. B: The network assortativity, a metric detailing the connectivity of nodes with similar degree, is assessed for the genome network (blue), a 5-edge rewiring (yellow), and a 10-edge rewiring (green). The graph shows that the assortativity of the genome network intersects the x-axis at two points-the first point occurring as the cells are transitioning into the neurosphere stage, while the second point occurs as the cells are transitioning beyond the neurosphere stage. C: 3D graph illustrating edge contribution of each node (chromosome) throughout the differentiation time course. Edge contribution (z-axis) of each node (x-axis) dynamically oscillates over time (y-axis), indicating the potential for hub formation of select chromosomes at specific time points during cellular commitment. Interestingly, there are a myriad of behaviors for each plot ranging from a fairly uniform line (chromosome 20) to one that oscillates frequently (chromosome 10). D: Heatmaps illustrate preferential associations between nodes throughout the differentiation time course. Several node pairs (orange/white) appear to interact repeatedly in the heatmap of the compressed genome network. The specificity between node pairs is absent in the heatmap of the compressed random network; i.e. preferential node association is lost.

To determine whether the highly connected nodes preferentially interact with one another over the time course, we measured the assortativity metric of the genome metric compared to the rewired and control networks (Fig 3B and S6B Fig). The analysis demonstrates the genome network transitions between negative and positive assortativity over the differentiation time course, while the control networks remain negative over time. We suggest that transition in assortativity of the genome network is due to the heightened transcriptional synergy occurring among the highly connected nodes (chromosomes) during the process of cellular commitment. This entrainment may occur due to an increased instance of bidirectional edges between nodes during network construction. The overall behavior of the genome network and the 5-edge rewiring is similar, although the rewired graph transitions between negative and positive assortativity more frequently and is altered in magnitude. In addition to transitioning between negative and positive values more frequently, the 10-edge rewiring also traverses the x-axis at earlier time points relative to its counterparts (Fig 3B).

As a means to ascertain which nodes may function as hub-like domains over cellular differentiation, we visualized the edge contribution of each node (chromosome) as a ratio of its degree to the total number of edges within the network over all time points (Fig 3C). On average, each chromosome contributes between 5-12% of edges present in the emergent genome network over the differentiation time course. The node profile for each chromosome varies over the time course and illustrates that potential hub formation is transient and relies upon temporal co-regulation as a function of the mean SKL between all differentially expressed genes on interacting nodes. Finally, we were interested if certain chromosomes preferentially interacted over the entire differentiation time course. We compressed the adjacency matrix of the genome networks, representing chromosome associations at each time point of the differentiation, to a single heat map and compared it to a compressed adjacency matrix created using an Erdős-Rényi random network distribution (Fig 3D). We observed that certain chromosome/nodes preferentially interact at moderate to high frequency in the genome network (Fig 3D, left), compared to promiscuous associations of the control heat map (Fig 3D, right), illustrating non-random, expression-mediated organization.

In sum, these measurements highlight three interesting features of the *in silico* chromosome association genome networks. First, all of the topological descriptors indicate properties substantially different from random networks; the genome networks possess a profile that is more “real-world” in nature (S5B Fig). Second, the specific connections of the nodes in the *in silico* chromosome association genome networks are highly sensitive to perturbation. Rewiring of the nodes of the genome network in a limited fashion leads to marked differences in the clustering potential and assortative capacity (Fig 3A, B), indicating that organization plays a pivotal role in maintaining its unique topology. Finally, transition in assortativity seems to be a unique feature of chromosome organization occurring during differentiation (Fig 3B and S6B Fig). This result suggests that similarity in gene co-regulation facilitates association of nodes of similar degree distribution, creating hub-like expression domains that drive lineage commitment.

### In situ analysis supports the validity of in silico predictive networks

In order to test our *in silico* modeling, we performed the hESC to cNPC differentiation strategy that yielded the gene expression data (S7A Fig) [39]. We confirmed expression of genes that are hallmarks of the differentiation at both the transcript and protein levels, using qRT-PCR and immunofluorescence, respectively (S7B Fig). Our results indicate the elevation of transcripts for *SOX1* and *COL3A1*, both associated with brain development, over the differentiation time course. We also detected a decrease in OCT4 at both the transcript and protein levels, indicating a loss of pluripotency upon commitment. The nuclear intermediate filament protein LMNA/C, which has been reported to be upregulated upon commitment of hESCs, was also elevated at the protein level while LMNB, a universally expressed nuclear intermediate protein, was present during both time points (S7C Fig). Additionally, small neurite-like protrusions emanating from the cell body became apparent at the end of the time course. These results indicate that we were successful in inducing formation of cNPCs.

To experimentally validate our *in silico* model of expression based genome organization, we visualized the pairwise association of eight chromosomes (HSAs) utilizing 3D-FISH at three points along the differentiation time course. The chromosome pairs were chosen based upon potential interactions at specific times within the context of our chromosome interaction genome networks (Fig 3B-D). Three chromosome pairs served as experimental conditions, exhibiting bi-directionality early upon edge addition during network construction at the indicated time points (S8A-C Fig): Day 0 HSA 7/HSA 20, 5 HSA 1/HSA 2, and Day 10 HSA 11/ HSA X. Two pairs, HSA 5/HSA 20 and HSA 1/HSA 3, lacked bi-directionality early during the edge addition process at any time point and served as controls (Fig 3B-D and S8A-C Fig). We predicted node bi-directionality of the experimental chromosome pairs would serve as a predictor for increased heterolog proximity at points of interest due to their transcriptional synergy. The single directionality and lack of graph neighborhood association exhibited by the control node/chromosome pairs were predicted to have reduced heterolog proximity at any time point due to asymmetry in transcriptional synergy.

3D FISH was performed with the pairs of chromosomes indicated above simultaneously visualized with fluorescently labeled whole chromosome paints (WCPs) at Days 0, 5, and 10 of the differentiation (Materials and Methods, 3D FISH). Confocal stacked images of the hybridized cells were captured and then converted into binary masks before being reconstructed into 3D renderings using the MATLAB Image Processing Toolbox (Fig 4A). To test our ability to predict chromosome organization as a function of co-regulated gene expression during differentiation we measured the clustering coefficient and percentage overlap of the heterolog pairs (Fig 4B). These methods of analysis were chosen as they address the salient features of chromosome organization in their ability to capture proximity (clustering coefficient) and physical association (overlap), both of which are important due to the amorphous nature of chromosome territories (CTs) (Fig 4A) and evidence suggesting that the territory periphery may harbor actively co-regulated genes [47, 48].

**Fig 4.**
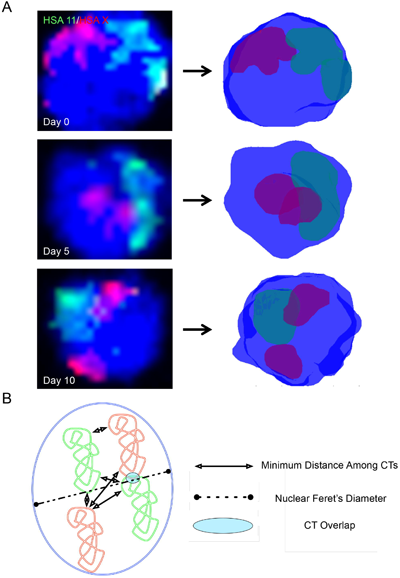
Analytic metrics for testing *in silico* modeling. A: The left column depicts representative 3D FISH analysis maximum intensity projections of DAPI stained nuclei (blue) hybridized with HSA 11 (green) and HSA X (red) at Day 0 (top), Day 5 (middle), and Day 10 (bottom). The right column depicts 3D renderings of the nuclei and chromosome territories of the left column. The software renderings were made by subjecting the thresholded masks of confocal Z-stacks at each time point using the smoothened isosurface algorithm in MATLAB. B: Cartoon schematic illustrating measurements between homologous and heterologous chromosomes. Signal overlap is measured as the percentage of chromosome overlap contributed by each territory on a per nucleus basis. The clustering coefficient is taken by calculating the mean minimum distance between all chromosome pairs as a ratio to the maximum nuclear (Feret’s) diameter.

The 3D clustering coefficient was calculated by taking the average pair-wise distance between the closest points of all chromosome pairs divided by the Feret’s diameter of the nuclear mask. This measurement theoretically gives an output ranging from 0 (closest) to 1 (farthest) as an indicator of the global proximity of chromosomes with respect to one another in the nuclear compartment. Comparison of the clustering coefficient of a given heterolog pair to that of all others allows us to determine the specificity of a decrease in minimum distance among all heterologs at a given time point in the differentiation. As stated above we hypothesize that minimization of distance between heterologs is predicted by the similarity of their composition of co-regulated genes during hESC to cNPC differentiation and thus their neighborhood association in our genome network modeling (Fig 2B-D).

We observed that the clustering coefficient of the HSA 1/HSA 2 and HSA 11/HSA X experimental pairs are significantly reduced at the predicted time points, Day 5 (HSA 1/HSA 2) and Day 10 (HSA 11/HSA X), respectively (Fig 5A). In the case of HSA 1/HSA 2, the minimization of distance between the heterologs was maintained on Day 10. While there was a trend of the HSA 7/HSA 20 pair to lose proximity on Day 5, it was not sustained and proved not to be significant (Fig 5A). The lack of significance in the association between HSA 7/HSA 20 does not align with our hypothesis; however, the global level of chromatin decondensation observed within hESC nuclei may render it difficult to model an expression-mediated chromosome organization paradigm during the state of pluripotency in contrast to the active reordering of gene expression during cellular differentiation [35]. As predicted by our model, there was no significant clustering of the HSA 1/HSA 3 control pair during any time point. Yet, there was a gradual reduction in the clustering coefficient of the HSA 5/HSA 20 heterolog pair, with a significant difference observed between Day 0 and Day 10. This unexpected result may be due to both HSA 5 and HSA 20 being present in node neighborhoods (Fig 2C, D) shared by at least one nucleolar organizing region (NOR) containing chromosome (HSAs 13, 14, 15, 21, and 22) at Days 5 and 10. NOR enriched chromosomes have been shown to influence genome organization, which may be accentuated due to size/structural changes occurring at nucleoli upon neuronal development [49-51].

**Fig 5.**
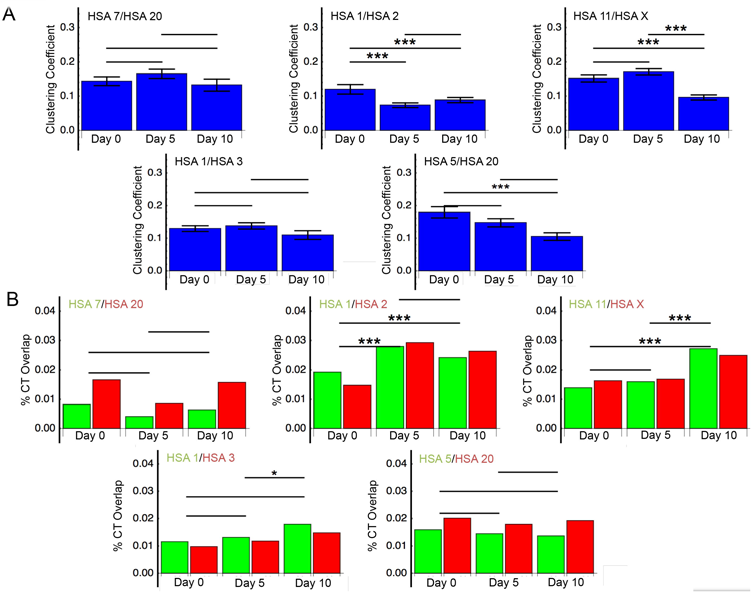
*In situ* validation of interchromosomal associations predicted *in silico*. A: Graphs detailing the clustering coefficient, calculated as the mean pairwise distance between all chromosomes as a function of maximum nuclear diameter, of experimental (HSA 7/ HSA 20, HSA 1/HSA 2, and HSA 11/HSA X) and control chromosome pairs (HSA 1/HSA 3, HSA 5/HSA 20) at Days 0, 5, and 10 of differentiation (26<n<102). *:p<=.05, **:p<=.01, ***:p<=.005; T-test. B: Graphs illustrating the signal overlap between experimental heterologous pairs (HSA 7/ HSA 20, HSA 1/HSA 2, and HSA 11/HSA X) and control heterologous pairs (HSA 1/HSA 3, HSA 5/HSA 20) at Days 0, 5, and 10 of the differentiation (26<n<102). Colors of text indicate the fluorophore that was used in the 3D FISH analysis (green=FITC, red=TxRed). *:p<=.05, **:p<=.01, ***:p<=.005; Mann Whitney Test.

While the clustering coefficient provides a metric for global heterolog proximity, to directly test whether the physical association of heterolog pairs is predicted by gene co-regulation during cellular differentiation, we measured the percentage of the overlapping regions of their CTs (Fig 5B). As indicated above this metric addresses that the periphery of chromosome territories may harbor actively co-regulated genes [47, 48], which would be facilitated by the association of heterologs with similar expression profiles. In support of the clustering coefficient analysis, we determined that the HSA 1/HSA 2 and HSA 11/HSA X experimental pairs demonstrate significant percentage CT overlap at the predicted time points, with the degree of overlap consistent with previous studies of functional chromosome associations (Fig 5B) [52]. Intriguingly, despite a trend of the HSA 7/HSA 20 pair to have heightened association at Day 0 (i.e. hESCs), the lack of a significant relationship supports the idea that the general state of decondensation accompanying pluripotency may impact the ability to predict expression-based chromosomal interaction networks. Unlike the clustering coefficient, the control two control heterolog pairs, HSA 1/HSA 3 and HSA 5/HSA 20, largely demonstrated no significant patterns of overlap. Therefore, while NOR containing chromosomes being present in node neighborhoods may influence global association of other nodes, they do not appear to impact the physical overlap of adjacent chromosome territories.

We also calculated the changes in homolog coalescence of the chromosomes analyzed above during hESC to cNPC differentiation. Decreased distance, particularly in the instance of coalescence, between homologous chromosomes may facilitate expression of gene subsets important at a specific differentiation time point. While our modeling approach was based on heterolog associations, our previous work showed an increased association of homologous chromosomes during murine hematopoiesis as a function of their coregulated gene sets [18]. We therefore sought to determine if there was also a correlation in homolog coalescence and cellular commitment of hESCs by quantifying the percentage of nuclei containing one discernable chromosome of interest at three time points during differentiation (S9A Fig). Similar to our previous findings, we observed an increase in homolog coalescence from Day 0 to Day 10 with all chromosomes except HSA 20 and HSA 2 (S9B Fig). While the degree (and therefore significance) of coalescence varied among the chromosomes, the trends are clear. Interestingly, both HSAs 20 and 2 demonstrate a pronounced level of coalescence at Day 0 (hESCs), such that an increase would be exceptional, and indeed reveal a reduction over the time course (S9B Fig). Regardless, these results indicate that homolog coalescence is also associated with hESC differentiation and may facilitate coordinate gene regulation.

Given the above caveats, we examined additional chromosome pairs to more robustly test our *in silico* model. Our analysis of HSA 7/HSA 20 suggests that the genomic state of pluritpotency may affect the role of heterolog association in coordinating gene expression during stem cell commitment. Thus, we analyzed the association of HSA 9/HSA 2, which also demonstrates bi-directionality early upon edge addition during network construction at Day 0 (Fig 2B-D and S8A-C Fig). In support of the influence pluripotency has on the ability to predict expression dependent heterolog associations, the pair reveals no significant difference among time points with either the GCC or CT overlap metrics (Fig 6A, B).

**Fig 6.**
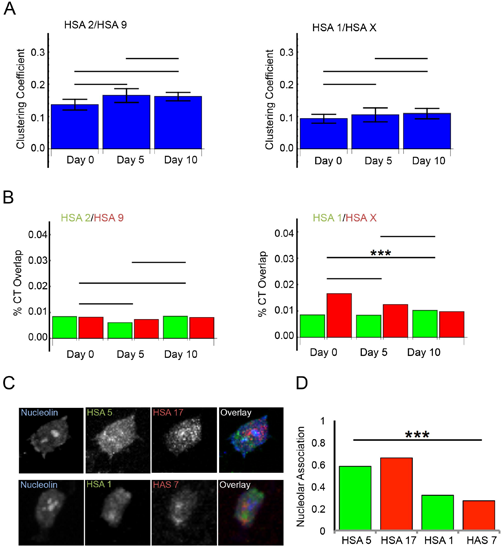
*In situ* validation of interchromosomal associations predicted *in silico*. A: Graphs detailing the GCC, calculated as in Fig 6, of a heterologous pair (HSA 9/HSA 2) that displays bi-directionality at Day 0 and a heterologous pair (HSA 1/HSA X) that demonstrates network neighborhood association at all time points of the differentiation (n≥50). *:p<=.05, **:p<=.01, ***:p<=.005; T-test. B: Graphs illustrating the signal overlap between heterologous pairs HSA 9/HSA 2 and HSA 1/HSA X, assayed as in Fig 6, at Days 0, 5, and 10 of the differentiation (n≥50). Colors of text indicate the fluorophore that was used in the 3D FISH analysis (green=FITC, red=TxRed). *:p<=.05, **:p<=.01, ***:p<=.005; Mann Whitney Test. C: Images representative 3D immunoFISH analysis maximum intensity projections of nucleolin (nucleoli) (blue) hybridized with HSA 5 (green) and HSA 17 (red) top column, and HSA 1 (green) and HSA X (red) bottom column. This analysis was performed at Day 5 of the differentiation, when HSAs 5 and 17 associate in a network neighborhood with NOR-containing chromosomes, while HSAs 1 and X do not. at Day 0 (top), Day 5 (middle), and Day 10 (bottom). D: Quantification of the nucleolar association of heterolog pairs in (C). Projections were scored for whether either HSA in a given nuclei revealed overlap with the nucleolin stain (n≤50). *:p<=.05, **:p<=.01, ***:p<=.005; One-Way ANOVA. T-test evaluations between heteolog pairs 5/17 and 1/X were not significant, but all other pairing permutations were significantly different (p<=.0001) as anticipated.

We next tested our model using an alternative approach in selecting chromosome pairs to be analyzed. Instead of choosing heterologs based upon potential interactions at specific differentiation time points within the context of the chromosome interaction genome networks, we analyzed the HSA 1/HSA X pair which demonstrates bi-directional node association across all three time points (Fig 2B-D and S8A-C Fig). Thus, the model would predict a degree of proximity that should be shared at Days 0, 5, and 10. Indeed, using our analysis approach described above we find that the pair does not significantly change in regards to the GCC during differentiation (Fig 7B). CT overlap similarly shows little change between Days 0 and 5, and Days 5 and 10 (Fig 6B). However, there is a significant change in regards to Days 0 and 10. A closer look at the network neighborhood at Day 0 reveals that HSA 1 and HSA X are its exclusive members, whereas at Days 5 and 10 there are several more chromosomes participating in the neighborhood (Fig 2B-D and S8A-C Fig). Therefore, it may be relevant to consider the ‘size’ of the network neighborhood when considering the predictive capability of our model, as the number of available nodes (chromosomes) may influence the association of a given pair.

Finally, in our initial analysis we observed that the control pairs HSA 5/HSA 20 demonstrated unpredicted proximity in our clustering coefficient analysis at Days 5 and 10 (Fig 5A). Upon closer examination of their node networks, we recognized that NOR-containing chromosomes make up numerous connections at those time points, leading us to speculate that NOR chromosomes may affect at least the clustering coefficient metric (Fig 2C, D). Therefore, to test this possibility we chose two pairs, HSA 5/HSA 17 and HSA 1/HSA 7, which have multiple NORs in their node at Day 5 and none, respectively (Fig 2C and S8C Fig). We performed 3D immuno-FISH with an antibody to nucleolin and relevant chromosome paints to identify the nucleolus and measure the association of the CTs with it. In support of the importance of NORs in predicting chromosomal associations as a function of their association at nodes, we found a significant enrichment for nucleolar association with the HSA 5/HSA 17 pair compared to HSA 1/HSA 7 (Fig 6C, D). Therefore, as with pluripotency, it is important to consider NORs when predicting chromosomal associations based on network interconnectivity.

## Discussion

In our study we have formulated an *in silico* model describing how similarity in the expression profile of differentially regulated gene networks is correlated with the higher-order organization of chromosomes during differentiation from hESCs to cNPCs. The refinement of gene expression patterns over the duration of cellular commitment is well documented [52-54]. Our results highlight that entrainment of co-regulated gene expression during differentiation is a dynamic process (Fig 1). Using graph statistics we observe that the genome networks generated using our *in silico* model exhibit properties similar to real-world networks (Figs 2, 3 and S5 Fig). We tested our modeling by assaying relative nuclear organization using 3D FISH of several chromosome pairs predicted to be proximal (Figs 4-6). While previous reports have related the positioning of chromosomes within the nucleus to size and gene density, our results support our earlier findings that relative proximity of the chromosome complement is correlated with the transcriptional synergy of differentially expressed genes on adjacent chromosomes during interphase [18, 19]. Our *in silico* modeling approach provides the ability to assess for potential genome-wide interchromosomal associations that will facilitate directed 3C-derived strategies.

Although we were unable to construct complete diploid genome networks due to the inherent inability to assign genome-wide transcript levels to homologs, we validated our hypothesis of an expression-mediated organizational paradigm by constructing haploid *in silico* networks based upon ranked mean SKL distance. Analysis of degree distribution across nodes of the *in silico* model indicates a deviation from normality, a common feature of biological networks (S5B Fig) [44, 45]. We further characterized the genome networks using the metrics of GCC and assortativity, identifying their uniqueness relative to both control networks (Erdős-Rényi random and Barabási-Albert scale-free) (S6A, B Fig) and during their own rewiring (Fig 3A, B). In essence, these analyses suggest that a small number of chromosomes (nodes) in a given modeled network neighborhood may serve as functional hubs in which genes expressed at similar levels from different chromosomes may localize to facilitate the regulation of transcription, for example by associating with transcription factories. Regardless, visualizing edge contributions of the various nodes suggests that hub potential varies dynamically throughout neurogenesis.

To validate our model we analyzed the pair-wise 3D nuclear organization of eight chromosomes chosen for their predictive or control value. The results with two experimental pairs, HSA 1/HSA 2 and HSA 11/HSA X, confirmed our hypothesis of increased proximity at the expected differentiation time points (Fig 5). However, the third experimental pair, HSA 7/HSA 20, while demonstrating a trend for proximity at Day 0 (hESCs) as anticipated, did not reveal a significant relationship. This result suggests that pluripotency is an important consideration when attempting to model gene networks and their role in the higher-order organization of chromosomes during differentiation. As discussed above, pluripotency is related to a unique decondensed chromatin state that may make it difficult to predict expression-based patterns of genome organization. To further test this possibility, we chose an additional homolog pair (HSA 2/HSA 9) with expected proximity in hESCs (Day 0). In support of our inference, this pair also did not demonstrate a significant trend at the predicted time point (Fig 6A, B). Thus, additional information may be necessary in training our *in silico* generated genome network to accommodate pluripotency, such as the change in overall volume of chromosome territories as a function of differentiation.

While our control chromosome pairs, which were chosen due to their presence in disparate node neighborhoods (Fig 2B-D and S8 Fig), largely conformed to our hypothesis, there was an exception with the GCC metric of HSA 5/HSA 20. This pair demonstrated significant proximity during the differentiation time course at Days 0 and 10. Close examination of the interaction networks of HSAs 5 and 20 revealed that at these time points, both share node neighborhoods with at least one NOR-containing chromosome (HSAs 13, 14, 15, 21, and 22). These chromosomes have been shown to influence genome organization, which may be accentuated during neuronal development [49-51]. To directly test this possibility, we identified two chromosome pairs at Day 5 that are involved in node neighborhoods with or without NOR-containing chromosomes, HSA 5/HSA 17 and HSA 1/HSA 7, respectively, and performed immunoFISH (Fig 2C and Fig 6C, D). Our analysis corroborates the influence NOR-containing chromosomes have on chromosome proximity as HSAs 5 and 17 both demonstrate significant association with the nucleolus as compared to the controls. Therefore, as with pluripotency, modeling expression-based genome organization must also take into account structural features of the nucleus, in particular the nucleolus, that while not directly related to the expression network of a given state of differentiation may nonetheless influence chromosome adjacency patterns by the attendant needs of cell proliferation and function.

*In situ* analysis of genomic organization using whole chromosome paints documents the myriad changes occurring during differentiation of hESCs. In addition to our analysis of *in silico* predicted heterolog proximity, we found that homologs generally demonstrate the tendency to coalesce at increased frequencies through the differentiation time course (S9 Fig). We have previously observed this trend in murine hematopoietic differentiation, suggesting that this form of organization may be a common developmental feature [18, 19]. Although why this may occur has yet to be fully explained, we have evidence suggesting that homologous gene loci that are spatially proximal in the nuclear compartment tend to have similar expression levels, i.e. reduced transcriptional noise [29]. It is possible that loci residing in large chromatin domains on chromosome homologs could potentially behave similarly. As co-regulated gene sets are proximally distributed along their chromosomes, association of homologs could lead to increased transcription of genes involved in cellular differentiation to a particular lineage [18].

As gene expression is inextricably linked with cellular state, investigating organizational paradigms that facilitate efficient execution of regulatory networks may prove to be critical in assessing the identity of individual cells within heterogeneous populations. Specifically, identification of defined genome arrangements in terminally differentiated cells through modeling may accelerate development of novel *ex vivo* differentiation strategies when used in conjunction with traditional cell characterization methods such as RNA-Seq and flow sorting. There is real therapeutic potential in such studies, as correcting damaged tissues with functionally reprogrammed progenitor cells continues to show promise in treating intractable disease states.

## Methods

### Cell culture

The NIH-approved human ESC cell line WA-09 (H9) was used to generate committed neural precursor cells. The cell line was cultured feeder-free according to WiCell Research Institute recommended procedures using Matrigel and TeSR1 media. Differentiated colonies were removed via aspiration after dispase treatment. Colonies were then scraped gently and grown in suspension media (1:1 DMEM/F12+l-glutamine and Neurobasal media, 1x N2 and B27 supplements, 20ng/ml insulin, 20ng/ml bFGF, 20ng/ml EGF) in non-tissue treated culture dishes for a period of 6 days, with intermittent 10ml changes of media every other day. Spheres were collected, gently triturated, and grown on polyornithine-coated (5ng/ml for 1hr at RT) tissue culture treated plates for an additional 4 days in expansion media (DMEM/F12+l-glutamine, 10% BIT 9500, 20ng/ml bFGF, 20ng/ml EGF, 2ug/ml heparin), with 10ml changes of media every other day.

### Immunocytochemistry

Human ESCs differentiated for 0, 5, and 10 days were fixed in fresh 4% formaldehyde/PBS++ for 10 minutes at room temperature. Cells were then washed in 1x PBS, permeabilized in .5% Triton X-100/PBS++ for 10 minutes, and blocked for 30 minutes in 4% BSA/PBS++. Primary antibodies against OCT4 (Santa Cruz, C-10), LMNA (Jol3, Abcam), LMNB (Santa Cruz, C-20), and MAP2 (L.I. Binder, AP14) were applied to cells (diluted according to manufacturers specifications in 4%BSA) for 1 hour at 37C before washing in .5% Triton X-100/PBS++ for 10 min and application of detection antibodies (diluted according to manufacturers specifications in 4%BSA) for an additional 45 minutes. Stained cells were then washed in .5% Triton X-100/PBS++ for 10 min and mounted with Prolong Diamond with DAPI before imaging on the Nikon A1R confocal microscope.

### Quantitative Real-time PCR

Human ESCs differentiated for 0, 5, and 10 days were harvested via centrifugation, washed with 1x PBS, and lysed in TriZol. Total RNA was reversed transcribed using the Superscript III First-Strand Synthesis System. 10ng of cDNA product was used in conjunction with LightCycler 480 SYBR Green I Master Mix to ascertain relative fold change difference with respect to OCT4 (F:TCAGGAGATATGCAAAGCAG, R:CACTGCAGGAACAAATTCTC), SOX1 (F: GACGTTCCCACATTCTTGTC, R:CACCGAAGTTCAGTCTAAAAC), COL3A1 (F:AAGAGTGGAGAATACTGGGT, R:AACTGAAAACCACCATCCAT), and GATA4(F:ACCCCAATCTCGATATGTTT, R:CCGTTCATCTTGTGGTAGAG) using the LightCycler 480. All genes were referenced to B2M (F:ACTTTGTCACAGCCCAGAT, R:GCATCTTCAAACCTCCATGA).

### Bioinformatics

Microarray data detailing the 10-day differentiation of H9 hESCs to committed neural precursor cells was provided by Alexey Terskikh [39]. The raw data was quantile normalized before redundant and obsolete entries were removed. Significance Analysis of Microarrays [40] was applied to the resulting dataset to identify differentially expressed genes over the entire 10-day time course.

Genomic order was assessed by calculating the relationship between all differentially expressed genes across the genomic complement by using the Symmetrized Kullback-Leibler (SKL) divergence equation,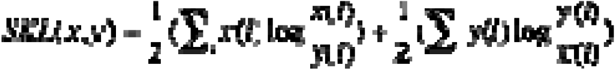, where x(i) and y(i) represent the normalized expression score of any two differentially expressed genes at a given time point. At each time point the resultant 2040×2040 distance matrices were hierarchically clustered with n=4 clusters. Genomic order is defined as the number of constituents in the largest cluster divided by the total number of differentially expressed genes.

Genome networks were constructed by creating matrix, M, detailing the mean SKL of the differentially expressed genes of each chromosome to all other chromosomes. The distance matrix, N, was normalized such that 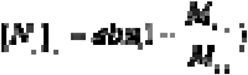. Each column of N was sorted in ascending order, with the highest association rank being 0, occurring between homologous chromosomes along the diagonal. Edges were added between nodes based upon the sorted rankings until a unified graph was formed, with one edge per node added at a time.

Given a graph 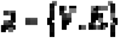, the assortativity is calculated by partitioning the vertex set V into k subsets such that each subset belongs to one community. The community modularity Q of this partition is defined as 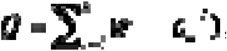, where e_ii_ is the percentage of edges that have both ends in community V_i_, and a_i_ is the percentage of edges that start from community Vi. In other words,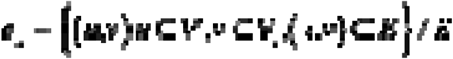 and 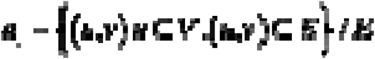. The graph metrics, i.e. the global clustering coefficient and the assortativity, were calculated with native Mathematica functions using the unified graphs.

The ER random networks and scale-free networks were constructed using the native Mathematica functions RandomGraph[n,k] and RandomGraph[BarabasiAlbertGraphDistribution[n,k]], respectively, where n is equal to the number of nodes (23) and k is equal to the number of edges present in the cohesive genome network at each time point. In order to mimic the behavior of the genome networks, self-loops were allowed to form during the construction of the control networks.

Chromosome association was analyzed using the MATLAB 2014a image processing toolbox. To determine distance measurements and overlap percentage among chromosome pairs within the nucleus, we used the actual voxel information of the binary masks (representing the chromosomes) that were created. In the case of non-normal distributions, Mann-Whitney tests were used to compare distance measurements between populations. Otherwise, in the case of normality, the T-test was used.

All scripts used within this section were executed in MATLAB 2014a and/or Mathematica 10.

### 3D FISH

Human ESCs differentiated for 0, 5, and 10 days were treated with Accutase for 10min at 37C and harvested via centrifugation at 1100rpm. Cells were resuspended in PBS and plated on poly-l-lysine coated coverslips before fixation in 4% formaldehyde for 10min. Cells were permeabilized with .5% Triton X-100 for 10 min, washed with PBS++ for 10 min, and incubated in .1M HCl for 5min. The slides were then washed in 2x SSC for 5 min before incubating overnight in 1:1 4x SSC/Formamide. Chromosome paints (MetaSystems) were applied to the cells and hybridized at 75C for 5min before storage at 37C for 2 days. Paints were removed by a 3x-wash in 2x SSC at 37C at 10min each wash followed by a 3x-wash in .1x SSC at 60C at 10min each wash. Slides were mounted using Prolong Diamond with DAPI and imaged on a Nikon A1R confocal microscope.

### 3D ImmunoFISH

The protocol above was performed with additional steps to incorporate the detection of nucleoli with an antibody to nucleolin, a canonical nucleolar protein. In short, after FISH preparation, rubber cement was removed, and cells were washed in 2X SSC three times for 5 min each at 37°C with gentle shaking and then 0.1X SSC at 60°C three times for 5 min each with gentle shaking. Cells were then rinsed in 4X SSC/0.2% Tween-20 in PBS, and then blocked in 4X SSC/0.2% Tween-20/4% BSA for 45 min at 37°C. Cells were then incubated with antibodies diluted in 4X SSC/0.2% Tween-20/1% BSA for 45 min at 37°C. Preparations were then mounted and imaged as above.

## Acknowledgements

Imaging work was performed at the Northwestern University Center for Advanced Microscopy generously supported by NCI CCSG P30 CA060553 awarded to the Robert H Lurie Comprehensive Cancer Center. We also thank Chelsee Strojny-Okyere for critical technical assistance. This study was funded by NIH grant DP2 OD008717 to S.K.

## Author Contributions

K.L. and S.T.K. designed research; A.G.G.G. performed experiment related to Fig 7C, D; E.K. and B.D. performed differentiation related to Fig 7A, B; A.T. provided expression data, K.L. performed research; K.L. and S.T.K. analyzed data; and K.L, E.S., and S.T.K wrote the manuscript.

## Conflict of Interest

The authors declare no conflict of interest.

## Supporting Information

**S1 Table. Chromosome characteristics and co-regulated gene distribution.**

**S2 Fig. Linear regression analysis of genome-wide co-regulated gene distribution**

On a genome-wide level, the greater number of differentially expressed genes on a given chromosome has a strong correlation with its number of significant bins (or domains) (r=.629) from the Exact Binomial Test of the sliding window analysis of Fig 1.

**S3 Table. Mean k-means cluster trends and standard deviation**

**S4 Fig. Analysis of genomic order in hESCs undergoing mesodermal differentiation**

The analysis was performed as described for hESC to cNPC differentiation. Data was acquired from: Piccini, I., Araúzo-Bravo, M., Seebohm, G., and Greber, B. (2016). “Functional high-resolution time-course expression analysis of human embryonic stem cells undergoing cardiac induction.” <u>Genomics Data</u> **10**: 71-74.

**S5 Fig. Correlation of inherent properties of chromosomes and network edge distributions**

A: The community number, based on graph modularity, is correlated to the linkage density, the average number of edges per node, of the genome network at each individual time point during differentiation. Significance is assessed using the Pearson’s correlation coefficient.

B: The degree distribution of the genome network at each time-point is compared to an Erdos-Renyi random network possessing the same number of edges over each time-point of neurogenesis. In nearly all cases the degree distribution of the genome network exhibits a departure from normality.

**S6 Fig. Global clustering coefficient and network assortativity of genome network relative to Erd**ő**s-Rényi and Barabási-Albert networks**

A: The GCC of the genome network is increased with respect to both the Barabási-Albert network and the Erdős-Rényi random network. The high GCC of the genome networks imply that it possesses a graph topology similar to real-world networks.

B: When compared to the Barabási-Albert network and the Erdős-Rényi random network, the genome network exhibits a transition from a parabolic transition from negative to positive assortativity over time that may reflect dynamic changes in *in situ* interchromosomal association during cellular differentiation.

**S7 Fig. Validation of neurogenesis system through qRT-PCR and immunofluorescence.**

A: Schema illustrating cell morphology and molecular markers present at the indicated time points of the differentiation.

B: Quantitative RT-PCR demonstrating the relative fold-change of transcripts involved in the maintenance of pluripotency (OCT-4), neurogenesis (SOX1 and COL3A1), and myogenesis (GATA4).

C: Immuno-staining of human embryonic stem cells (hESCs) and and committed neuronal precursor cells (cNPCs) using antibodies specific for nuclear markers (LMNA/C, LMNB), proteins involved in the maintenance of pluripotency (Oct4), and terminally differentiated neurons (MAP2).

**S8 Fig. Directional network used to choose whole chromosome probes for *in situ* validation of genome networks.**

A-C: Whole chromosome paints were chosen based upon node bi-directionality of the cohesive graph, indicated by red arrows, of edges during network construction at Day 0 (A), Day 5 (B), and Day 10 (C).

**S9 Fig: Homolog coalescence increased during cellular differentiation**

A: Cartoon diagram illustrating separate homologs compared to indistinguishable, coalesced homologs.

B: Coalescence frequency of each homolog pair used in the *in situ* validation experiments. In all cases, aside from HSA2 and HSA20, a gradual increase in homolog coalescence is observed over the 10-day differentiation time course. Bar graph colors indicate the fluorescence channels in which FISH was performed. Significance was determined by conducting a Fisher’s Exact Test in a pairwise fashion between time points: * p<=.05, **:p<=.01, ***:p<=.005; Fisher’s Exact-test.

